# 3D spatial organization and network-guided comparison of mutation profiles in Glioblastoma reveals similarities across patients

**DOI:** 10.1101/523563

**Authors:** Cansu Dincer, Tugba Kaya, Ozlem Keskin, Attila Gursoy, Nurcan Tuncbag

## Abstract

Mutation profiles of Glioblastoma (GBM) tumors are very heterogeneous which is the main challenge in the interpretation of the effects of mutations in disease. Additionally, the impact of the mutations is not uniform across the proteins and protein-protein interactions. The pathway level representation of the mutations is very limited. In this work, we approach these challenges through a systems level perspective in which we analyze how the mutations in GBM tumors are distributed in protein structures/interfaces and how they are organized at the network level. Our results show that out of 14644 mutations, 4392 have structural information and ~13% of them form spatial patches. Despite a small portion of all mutations, 3D patches partially decrease the heterogeneity across the patients. Hub proteins adapt multiple patches of mutations usually with a very large one and connects mutations in multiple binding sites through the core of the protein. We reconstructed patient specific networks for 290 GBM tumors. Network-guided analysis of mutations completes the interaction components that mutated proteins potentially affect, and groups the patients according to the reconstructed networks. As a result, we found 4 tumor clusters that overcome the heterogeneity in mutation profiles, and reveal predominant pathways in each group. Additionally, the network-based similarity analysis shows that each group of patients carries a set of signature 3D mutation patches. We believe that this study provides another perspective to the analysis of mutation effects and a good training towards the network-guided precision medicine.

**Author Summary:** Glioblastoma (GBM) is the most aggressive brain tumor type with a 15 months of survival on average. The mutation distribution of the GBM patients is very heterogeneous and standard treatments fail to consider the inter-tumor heterogeneity. In our study, we follow a systems level approach that integrates mutation profiles with protein-protein interaction networks. We hypothesized that although the mutations are heterogeneous, the mutations that are close in 3D of the same protein may affect the protein function similarly and this information can be used to get meaningful relation between disease state and mutations. Therefore, we spatially grouped these mutations as “patches” and reconstructed patient specific protein interaction networks. When we cluster these networks based on their pathway similarities, we found four patient groups in GBM. Then, the comparison of groups revealed overrepresentation of 3D patches. This finding can be used for patient specific therapy.

## Introduction

Cancer mostly occurs when somatic mutations accumulate and eventually change the behavior, structure and properties of the cell. Understanding which mutations cause cancer is of crucial importance. As a result of large-scale cancer genome sequencing projects including The Cancer Genome Atlas (TCGA) [1] and the International Cancer Genome Consortium (ICGC) [2] and smaller-scale gene/protein focused and genome-wide screenings have enabled to discover a large volume of somatic mutations in human cancers. However, not every somatic mutation causes cancer. The main challenge is to detect the driver and passenger mutations. While driver mutations provide positive growth advantages to cancer cells, passenger mutations do not provide selective advantages to them [3]. Ultimately, only the proteins from the driver genes are the molecular targets in drug discovery and cancer therapy. Also, having insights about the combination of mutations that cause cancer is equally important to understand the causes and mechanisms of cancer development and progression. Cancer is a heterogeneous disease. The diversity between and within tumors as well as among individuals of the same type of cancer is enormous. All these together with epigenetic and post-translational factors determine the risk of cancer progression and the therapeutic resistance. One drug that works in an individual might be ineffective in another individual. Therefore, more challenge occurs even when one tries to understand the driver mutations in a single cancer.

Proteins interact with each other to perform their functions in almost all biological processes including signaling, replication, cell to cell communication, transcription and many others. Alterations or mutations in proteins may affect their interaction, leading to dysfunction and disease. Interactome is the complete set of protein-protein interactions in an organism. Representing the full interactome may be impossible because of its complexity. Post-transcriptional modifications, cellular localization, tissue specificity further increase complexity. On the other hand, some proteins interact permanently, some proteins need phosphorylation to interact their partners and protein interactions could be tissue-or cell type-specific. This complexity makes it difficult to get a unique full interactome. Time complexity is another issue. PPIs are dynamic and are coordinated with respect to cell’s status. Experimental methods are insufficient to reveal the whole interactome for all the complexity, computational methods can also be used to complement/predict protein-protein interactions [4].

Several studies have focused on the impact of the disease-associated alterations in the protein-protein interaction networks [5–8]. Very recently, IMEX consortium [9] also started an effort to curate and catalogue the functional impact of mutations in protein interactions [10]. Combining three-dimensional structure data with large scale mutation data might help to elucidate the effect of mutations in cancer [6, 11–14]. A protein’s biological function and physical interactions are closely related to its structure. Mutations that completely disrupt a protein’s structure (i.e. that correspond to folding core) can have different functional consequences than mutations that do not affect the global fold but rather affect a surface or interface region. Given that a protein may have several interfaces, which interface is affected by the mutation will shed insights about the functional changes of that protein. Disease-associated mutations are more likely to affect protein interactions [15]. Different mutations in the same protein may lead to different interaction profiles and eventually different disease phenotypes [16–20]. The edgetic perturbations [15] in proteins thus need to integrate structural knowledge with biological networks to re-construct structurally enriched PPI networks [21–23]. Beyond the position of the mutations in sequence, their organization in 3D structures of proteins has been evaluated in cancer in several studies from different perspectives [5, 8, 24, 25]. Niu et al cluster the mutations from 19 different cancer-types and come up with druggable functional mutations [25]. Additionally, some other studies focus on patient-specific analysis of the molecular signatures in tumors [26–28]. The phosphoproteomic data from eight GBM patients have been previously used to demonstrate that network-guided comparison reveals commonalities and differences across patients [27]. In another network-based approach, mutations, transcriptional and phosphoproteomic data have been used to model patient-specific pathways in prostate cancer [28]. Network-based analysis was also used to distinguish the driver mutations from passenger mutations in GBM [29]. Very recently, the functional effects of mutations on protein interactions and signaling networks have been extensively reviewed in [14] from a view of biophysics.

Given the number of mutations deposited in TCGA, the number of structures in PDB and the number of known interactions between proteins, computational approaches are crucial for a system level analysis of the mutation effects on proteins, protein interactions and functional pathways in a patient-specific way. We applied a system level approach to the somatic missense mutations in 290 Glioblastoma (GBM) patients which is the most aggressive type of brain tumors. The mutation profiles are very heterogeneous across the patients and it does not track with the known transcriptional subtypes of GBMs or the known biomarkers such as IDH1 mutation. Given this heterogeneous profile of the GBM mutations, our main aim is to address the question that despite this heterogeneity, could we reveal some commonalities across patients when we investigate the 3D spatial arrangement of mutations in proteins and their organization in the patient specific protein-protein interaction sub-networks. In our approach, both the spatial arrangement of mutations in proteins and protein-protein interactions, and the patient-specific sub-networks inferred from mutation profiles are used to better classify GBM patients. We proceed in two directions: (i) finding the spatial arrangement of the mutations in the sub-networks and also across patients and (ii) reconstruction of the sub-networks that are primarily affected by the given mutation set (see Fig 1).

**Fig 1.**
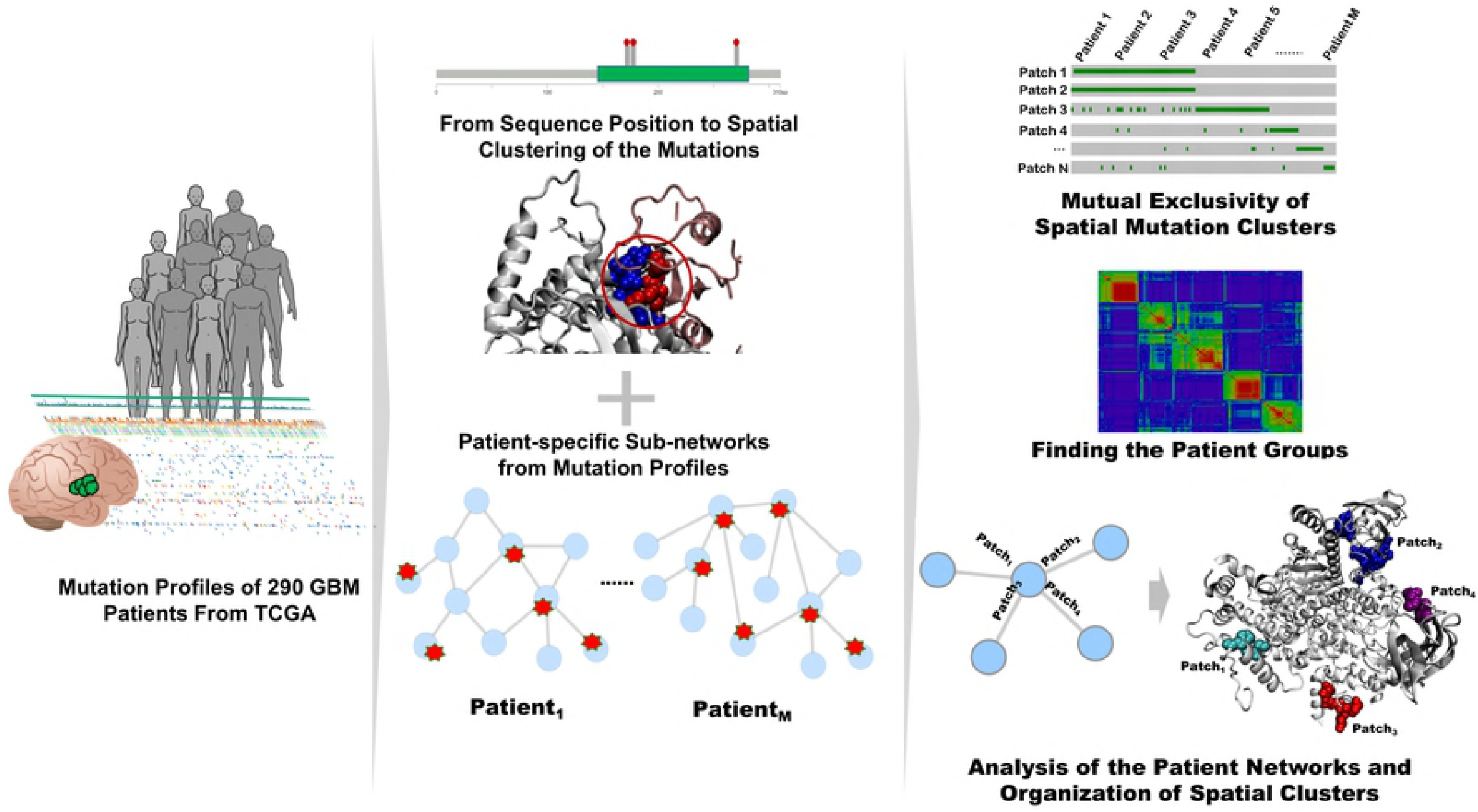
Overview of the method. Patient specific profiles are obtained from TCGA. Red dots and stars in the middle panel correspond to mutations mapped to the sequence, structure and PPI network (The icons in the first panel are retrieved from Reactome Icon Library [30]).

## Results

Missense mutation profiles are very heterogeneous across GBM patients at the sequence level. In this section, we present the findings of a system level approach to the somatic missense mutations in tumors from 290 Glioblastoma (GBM) patients.

### Heterogeneity partially decreases with 3D clustering of GBM mutations

The missense mutations from 290 GBM tumors were first analyzed at the sequence level. There are 14,644 unique mutations and 14,200 of them match at least to one protein. The average number of missense mutations per patient is 49.01. The distribution of the mutations across different GBM tumors is heterogeneous. Only 62 mutations are present in at least three patients and 245 mutations are present in at least two patients. The most frequent mutation with 18 patients is EGFR mutation A289V which is followed by IDH1 mutation R132H in 13 patients.

We mapped the GBM mutations onto protein structures and their local organization was determined by their spatial proximity which we call “patches”. Out of 14,200 mutations, 4392 mutations map to at least one protein structure either from PDB [31] or from ModBase [32]. A patch is a set of mutated residues such that either a mutated residue is physically in contact with another mutated residue (that is, at least one pair of atoms within 5Å distance), or there is another intermediate residue connecting two mutated residues. The term “patch” was used in previous studies, however we have to indicate that our patch definition is different than those [12, 25], because instead of using a mutated residue as a centroid of the patch, we search for continuous contacts of the residues. The grouping of 4392 mutations resulted in 220 patches and 3812 singletons (a mutation that cannot be put in a group). The 220 patches consists of 580 mutations. Each patch is found in 3.5 patients on the average, and each mutation is found in 1.13 patients on the average. The distribution of patches is different from the distribution of individual mutations as shown in Fig 2. The columns correspond to individual patients in both parts of the figure. We compared the presence of the mutations located in the patches individually (Fig 2A) and the presence of the patches in the patients with a frequency of at least 2% (Fig 2B).

**Fig 2.**
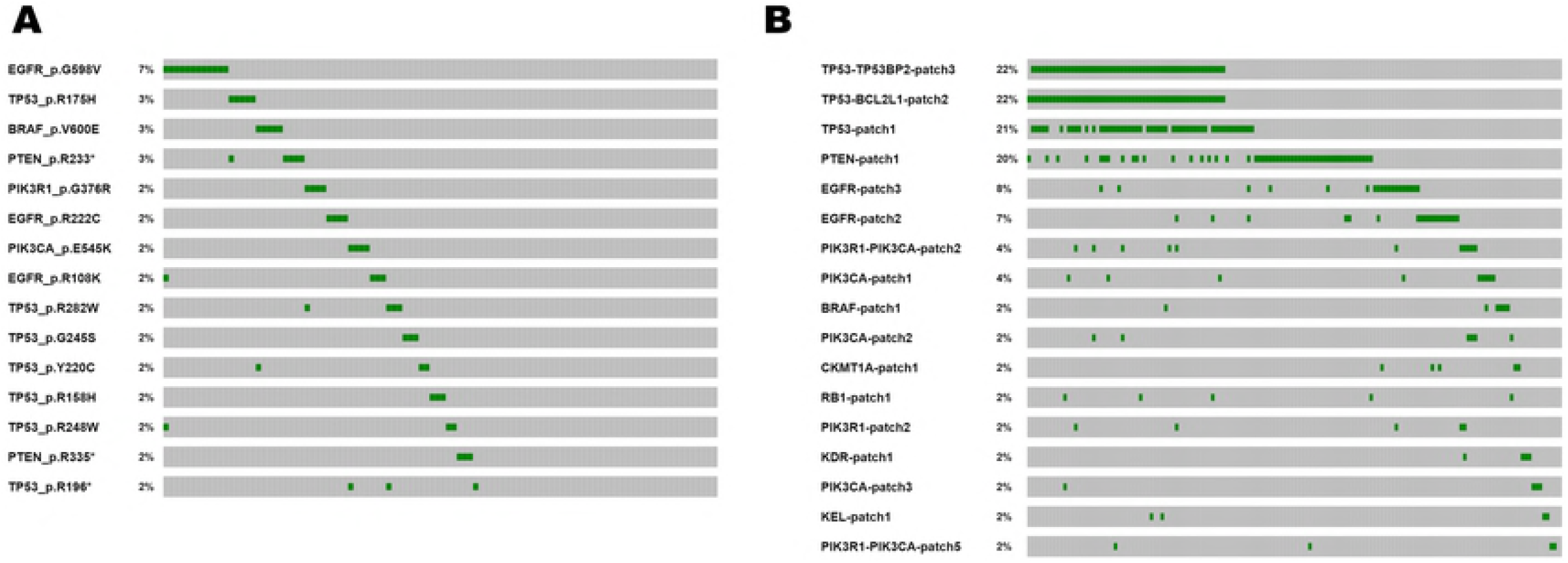
Single position versus 3D patch comparison of the mutations. (A) Mutations at a frequency of more than 2% of all patients. (B) Patches at a frequency of more than 2% of all patients. The Oncoprinter tool embedded in cBioPortal [33] was used to generate these oncoprints.

The majority of the patches contain two or three residues (Fig 3). These small patches tend to be intra-patch that is no residues from an interface region. The bigger patches happen to be interpatches (with at least one interface residue) and very large ones in the central proteins (TP53 with 41, PTEN with 43 residues as shown in the patch, Fig 3A). In total, there are 160 intra-, 60 interpatches in our dataset. An example of very large patches in PTEN is illustrated in Fig 3B. Some other examples of the patches are shown in Figs 3C-3E for the two patches of EGFR, the three patches of PI3KR1-PI3KCA complex where complementary chains have mutations and the interface patch of the SMYD2-TP53 complex, respectively. On the other hand, some large proteins, such as Titin, happen to have many mutations however, those mutations do not form a large patch but rather they are isolated (singletons). Earlier studies included Titin mutations as cancer related [34]. But later analyses considering the mutational heterogeneity carefully, identified TTN (and other large proteins such as olfactory receptors) to be false positives [35]. In our patch analysis, it is interesting to observe that Titin mutations to be isolated whereas in well-known cancer related proteins such P53, PTEN to have connected large patches. The patches in these major proteins seem to be in common among 20% of patients are in P53, PTEN and EGFR with 8%, overall yielding slightly better grouping of patients (see Fig 2B). 3D patches allowed us to cluster patients better in the same type of cancer. Previous studies suggest that 3D clustering of mutations led to better classification of different cancer types, driver mutations or novel cancer genes [8, 12, 25, 36]. Our results suggest that patient-grouping is also possible with 3D patches.

**Fig 3.**
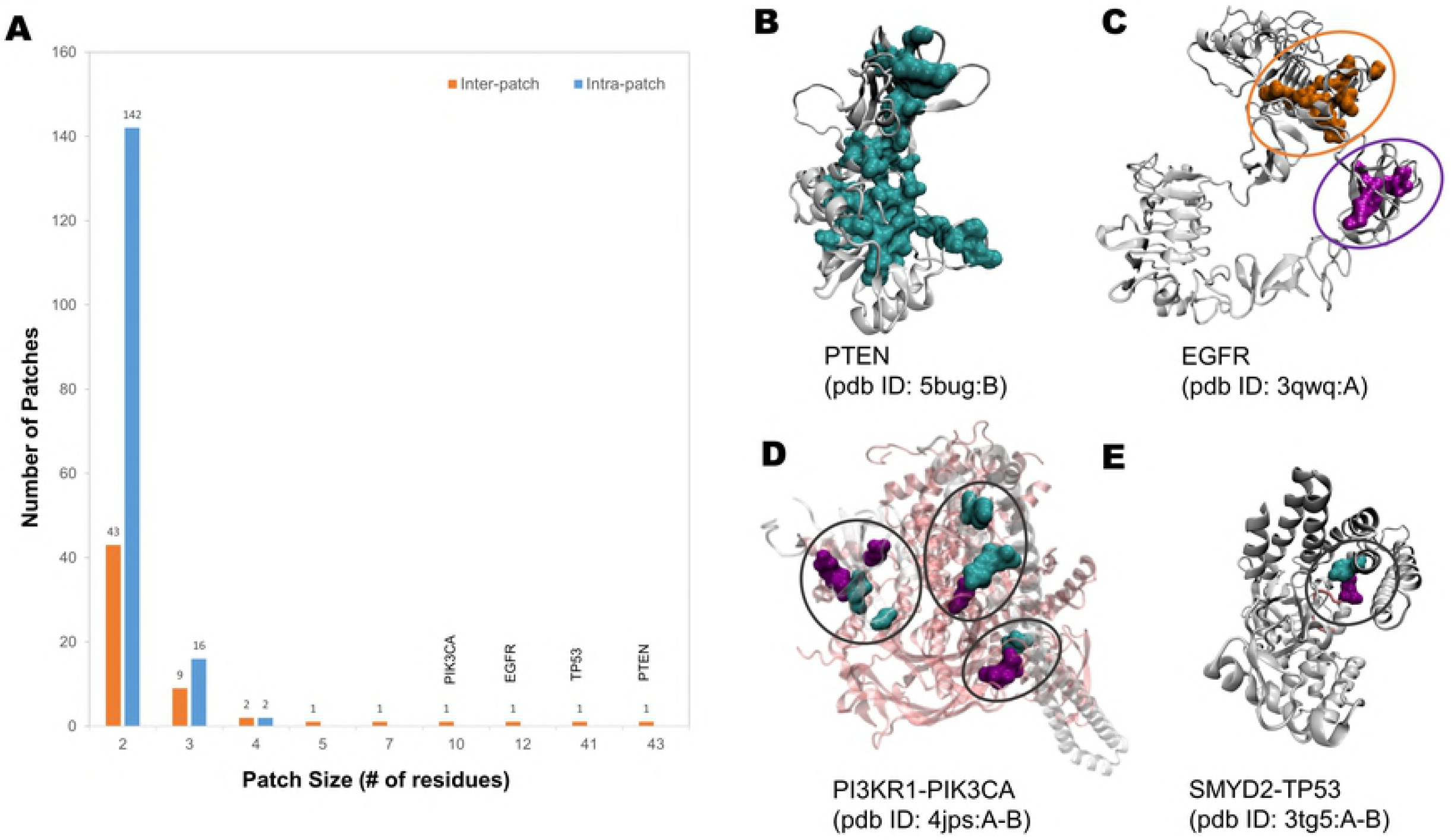
Patch size distribution and intra-/inter-patch examples. (A) Distribution of the patch sizes. (B-E) Examples of the patches in our dataset. (B) PTEN – patch size – 43 residues – an example of mutations forming a chain from the surface to core. (C) EGFR – two patches with different sizes, 10 and 7, respectively. (D) PI3KR1-PI3KCA – three patches – each patch has mutations in complementary chains. (E) SMYD2-TP53 – one patch with two mutations, each from one partner protein.

### Structural mapping of GBM mutations and physicochemical organization in proteins and protein interactions

We analyzed the characteristics of the GBM mutations at the molecular level from a chemical and structural perspective. Mapping mutations to protein structures enabled us to analyze the distribution of mutations on structures (whether they are located at the core, surface or interface regions). Protein structures are categorized as core (residues with no solvent accessibility), interface (amino acids of a protein making physical contacts with the amino acids of a partner protein), and the rest is categorized as surface (composed of solvent accessible residues excluding interface residues). We divided the mutations as singletons and the ones in the patches. Then, we analyzed the disease association of the singleton mutations and the patches on different protein regions.

#### Location and physico-chemical properties of mutations

The knowledge of the location and physico-chemical properties of mutations improve our understanding about their functional impact in the cell. Out of 14644 mutations, 4392 mutations are mapped to at least one protein structure. Of these, 3601 of these are at the surface and core of the proteins. PDB structures provided the location of 1522 mutations whereas, MODBASE model structures enriched this number by 2079. Most of the mutations are located in the surface region of these proteins based on the known structural data (see Table 1 for the summary of all numbers). The binding site information comes from PDB (if there exist complex structures, or from models of Interactome3D [6], Interactome Insider [13], and PRISM [37] as shown in Table 1.

**Table 1.**
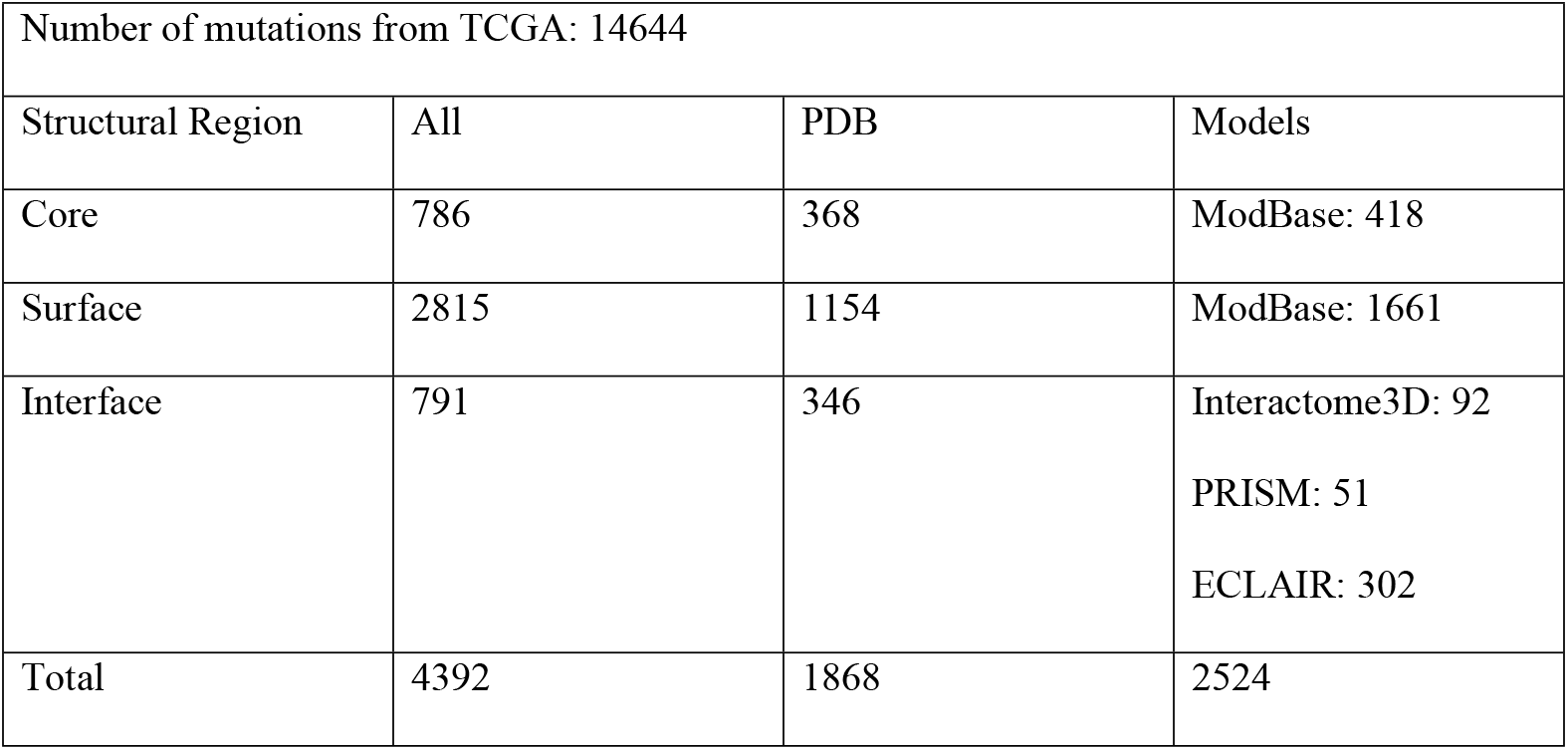
Number of mutations mapped to protein structures

We then checked what kind of alterations take place in chemical classes of the mutations. The chemical classes are described as hydrophobic, charged and polar. We analyzed whether a mutation conserves its wild type chemical class or it switches to another chemical class, such as from hydrophobic to polar (Fig 4A). A large portion of the core mutations is hydrophobic and their chemical class is preserved. Any change disturbing the hydrophobic characteristics of protein cores would most probably lead to the unfolded/unstable protein structures and would be lethal. Chemical class profiles of the interface and surface mutations are very similar to each other and substantially different from core mutations. The most prominent changes in surface and interface mutations are hydrophobic-to-hydrophobic, charged-to-charged and charged-to-polar shifts. We do not observe hydrophobic-to-polar and hydrophobic-to-charged changes in interfaces and surfaces either. These types of changes would most probably change the binding or solubility characteristics of the proteins.

**Fig 4.**
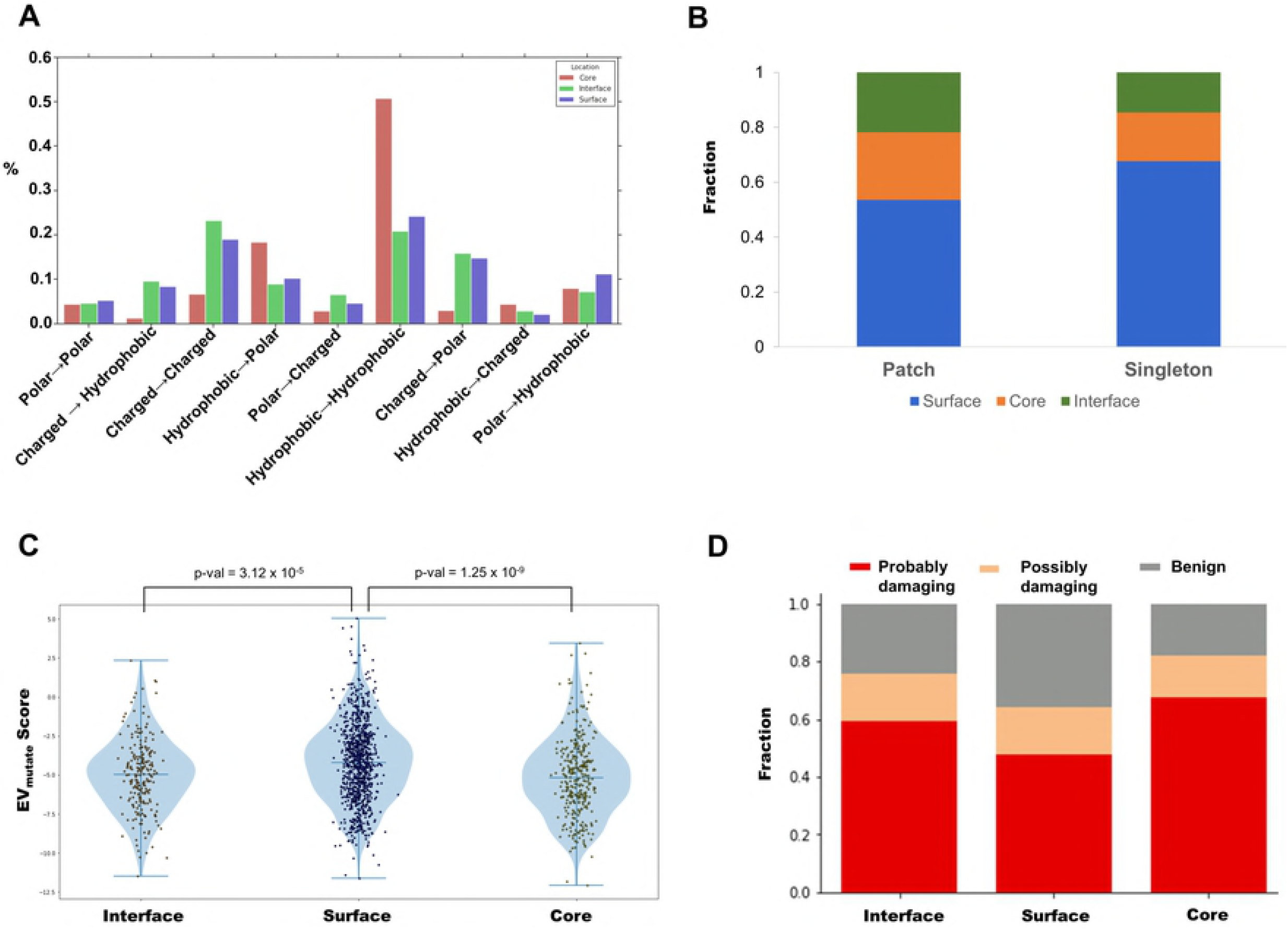
The characteristics of the mutations. (A) The changes in the chemical properties of the residues resulting from mutations according to their locations (B) Fraction of mutations that are organized in patch or stay as a singleton based on their location (interface, core, surface). (C) The distribution of the mutations according to their disease association in different locations according to EVmutation. (D) The fraction of the mutations according to their disease association in different locations according to PolyPhen-2.

Nishi et al found that GBM missense mutations on protein-protein interfaces have overall destabilizing effect and mostly alter the electrostatic component of binding energy [11]. They also showed that mutations on interfaces resulted in more drastic changes of amino acid physicochemical properties than mutations occurring outside the interfaces. David and Stenberg showed that there are differences in polarity, hydrophobicity and charge change between polymorphisms and disease causing mutations [38]. The distribution of physicochemical changes was significantly different between disease-causing SAVs and polymorphisms and in different regions of the proteins. Despite similar results, we further divided non-interface mutations into surface and core residues and found that interface properties are more similar to surface rather than the core mutations. This suggests that there might be interface regions over the surfaces that we do not know as of now.

Other interesting cases are that some positions are observed to be mutated differently in patients. There are 70 unique positions representing such cases. For example, Proline at 596th position of EGFR appeared to be mutated to Arginine in one patient while it is mutated to Leucine or Serine in other patients. We also checked their chemical property that 63% of all these positions are originally charged amino acids. The most frequent alterations in these positions are either preserving charged class or changing from charged to hydrophilic and polar. These are mostly found in interfaces and surfaces. Alterations to hydrophobic amino acids are very rare in these positions.

#### 3D mutation patches and disease association

Disease association of the mutations are not uniform across the proteins and their interactions. While most of the mutations prefer to be found as a singleton, approximately 13% (580 mutations) of the 4392 mutations are in close proximity to each other and form spatial patches. Additionally, out of 791 interface mutations, 145 are located in at least one patch (~18% of all interface mutations). When we compared the interface patch mutations to the non-interface patch mutations, we obtain an odds ratio of 1.63 (chi square P< 0.0001). Therefore, interface mutations prefer to be located at patches compared to non-interface mutations (Fig 4B).

We further assessed the effect of mutations for their disease-causing potentials using two different methods and analyzed those effects according to the location of the mutations. The first method is EVmutation [39] and the second is PolyPhen-2 [40]. Both methods gave similar results that mutations that are located in the core and in the interface are more damaging compared to the mutations located on the surface (Figs 4C and 4D), while the damage is more severe in the core region. The results also show that interface mutations organized in patches are more damaging compared to singleton ones. Similar results were obtained from EVmutation analysis. There are 60 patches having at least one interface residue.

We obtained PolyPhen results for 574 mutations located in the interfaces. There are 465 singletons (benign: 124, possibly: 79, probably: 262), and 109 mutations located in patches (benign: 15, possibly: 15, probably: 79) as shown in Table 2. 73% and 86% of singletons and patch mutations are possibly or probably damaging, respectively. However the behavior of hub proteins and others are vary from each other. When we separated our proteins as hubs (PTEN, TP53, EGFR, PIK3CA, RB1 and PIK3R1) and the rest, we observed that for the hub proteins, patch mutations are more damaging (98% of all patch mutations). There are too few singleton mutations that PolyPhen gave prediction results; therefore, we did not calculate the damaging ratio. For the rest of the proteins (excluding the hubs), however, 68% of patch mutations are damaging while 74% of singletons are damaging. In a previous study, we showed that energetically important hot spots can be found as singlets or clustered in hot-regions. Most of the disease causing single amino acid variations in that dataset-restricted to human proteins only and but not limited to cancer variations-were found as singletons rather than in hot-regions [19]. This result combined with the results here show that there can indeed be an evolutionary pressure on disease causing mutations to be clustered in the interfaces. Indeed, rare mutations are mostly found as singletons, since their damaging effect is more in the interfaces.

**Table 2.** Disease association of singleton and patch mutations in the hubs and the rest.

|  | Hubs |  | Rest |  | Total |  |
| --- | --- | --- | --- | --- | --- | --- |
|  | Patch | Singleton | Patch | Singleton | Patch | Singleton |
| Benign | 2 | 3 | 13 | 121 | 15 | 124 |
| Possibly | 9 | 0 | 6 | 79 | 15 | 79 |
| Probably | 58 | 1 | 21 | 261 | 79 | 262 |
| Total | 69 | 4 | 40 | 461 | 109 | 465 |

### Characteristics of interface mutations

Raimondi et al have analyzed somatic mutations in cancer through the protein interfaces and correlated the presence of some interfaces with the phenotype [41]. In the study of Meyer et al, besides introducing a new structural interactome, they also analyzed the organization of mutations in the protein interfaces [13].

As detailed in the previous section, 791 mutations were found to be in the interface regions of the proteins. Although it is a small portion compared to the total set of mutations, these residues have potential to affect 6283 interactions (See S1 File for detailed list for interface mutations and related interactions). When we further analyzed interface mutations we have seen that some mutations are distributed over the protein surface and located in multiple binding sites of the proteins that we call ‘multi-face’ mutations. In Fig 5A on the left panel, we illustrated an example to this type of mutations. PTPN11 interacts with GRB2 and ERBB2 proteins through different faces. Two mutations Q510L and E69K are located on different interfaces and the set of these mutations of PTPN11 can be classified as multi-face mutations.

**Fig 5.**
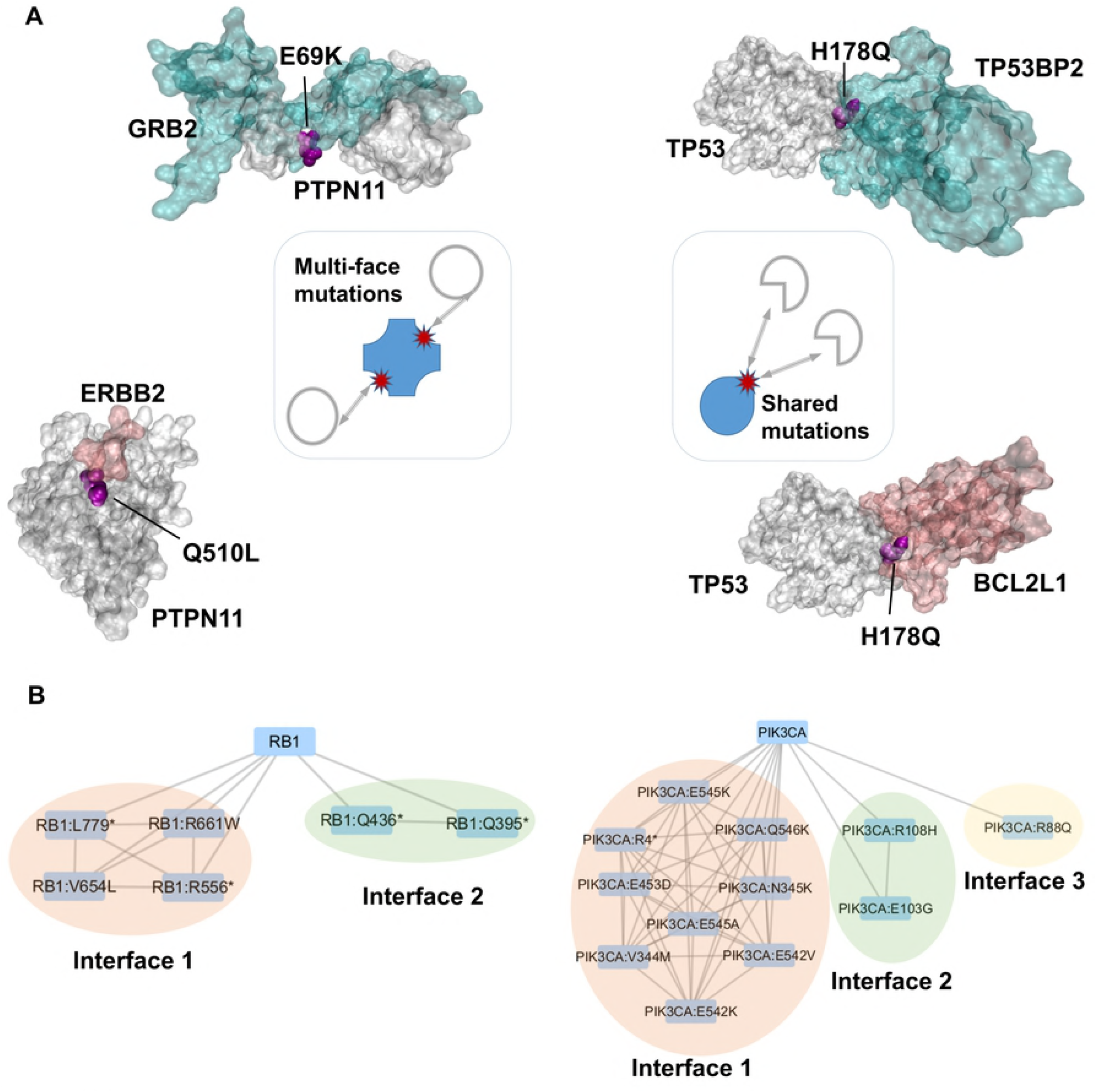
Representation of the multi-face mutations and shared mutations in interface. (A) Examples of both multi-face and shared mutations. Left part represents the examples for multi-face mutations and the right part represents the examples for shared mutations. (B) Organization of RB1 and PIK3CA interface mutations are represented as a network where the nodes are mutations and the edges represents at least one shared interaction between two mutations.

On the other hand, some mutations that are present in a binding site may affect multiple interactions because some proteins are adapted to use the same binding region to interact with multiple partners. We call this type ‘shared’ mutation. For example, the binding of TP53 to its partner proteins TP53BP2 and BCL2L1 occurs through overlapping regions and the mutation H178Q located in these interfaces is shared and may affect both interactions (see Fig 5). A network representation of these mutations are shown in Fig 5B where the edges between mutations represent that those mutations have shared binding partners. In RB1 protein, there are two binding regions and multiple mutations and in PIK3CA, there are three binding regions having mutations. In total, 211 proteins have only one mutation in their known binding site and only one protein binds through that region and from 211, only 12 are located in 3D patch which represents 6% of mutations. In the same class, 328 proteins have again only one mutation in their known binding site, but this site is repeatedly used by multiple partner proteins (shared mutations). These type of mutations are mostly present as singletons (only 6% are located in a 3D patch). Out of 791 mutations, 158 mutations are located in the different binding regions of 61 proteins (distributed in 105 different binding sites) and interact with multiple partner proteins. Among these mutations, 81 are located in a 3D patch (51%). Mostly, hub proteins such as PTEN, TP53, EGFR, are in this group. These hub proteins have multiple mutations and these mutations tend to be in 3D patches. 22 mutations belong to proteins that have multiple interface mutations (only 23% are located in a 3D patch) and interact with only one protein. On the other hand, 72 mutations again belong to proteins that have multiple interface mutations and interact with multiple proteins through the same site. Among these 72 mutations, 22% of them are located in 3D patch. In total, there are 505 shared mutations that appear in the same binding site and interact with many partner proteins through that region. These results show that shared mutations tend to be present as singleton. Moreover, the hub proteins have mostly multiface mutations which tend to be in 3D patches.

Yu and coworkers show that whether a mutation falls in known or predicted protein-protein interaction interfaces is related to the likelihood to disrupt these interactions, and mostly these mutations are found on the interfaces of hub proteins [42]. Overall, they suggest that network topology should be considered when interpreting the impact of mutations. Sahni et al also states that integrative analysis of mutations with protein networks can improve our understanding about the phenotype-genotype relations in diseases [15]. The results indicate that highly connected hub proteins, prefer having multiple patches in the interface regions and interface mutations are mostly located in the patches in the hubs. Proteins having multiple patches in their interface regions are interactome hubs and they are TP53, EGFR, PTEN, PIK3CA as shown in Figs 6A and 6B.

**Fig 6.**
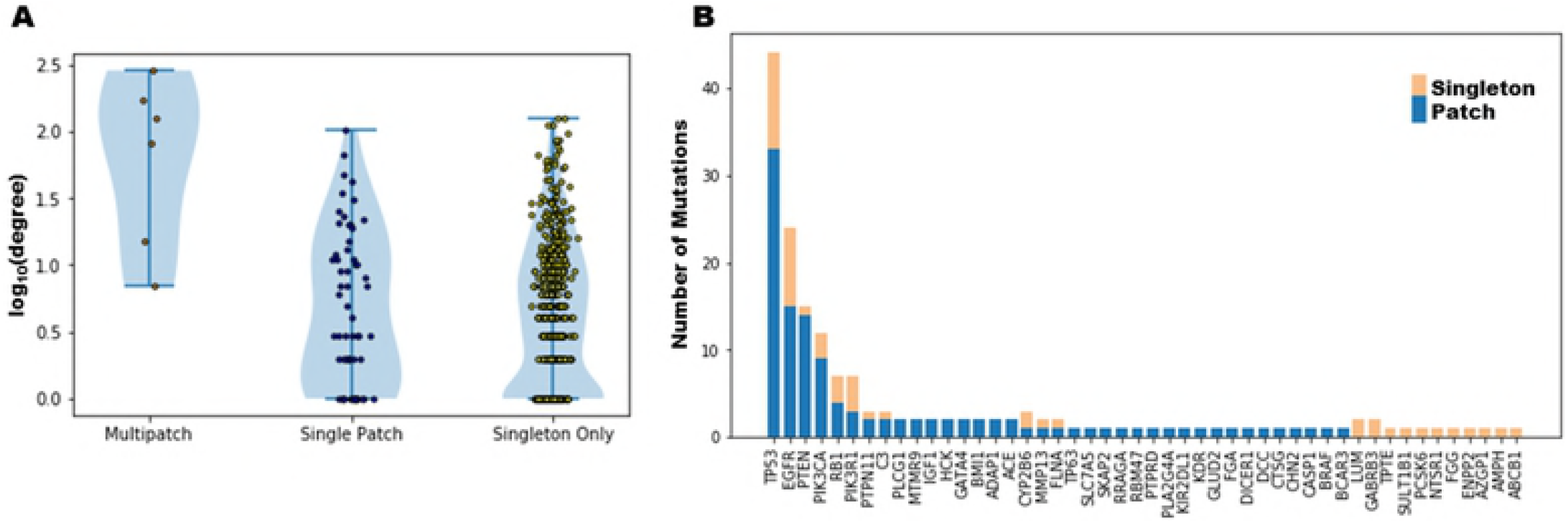
Relationship of the proteins having interface mutations and preferences about the spatial arrangement of these mutations. (A) High degree proteins (hubs) in interactome have tendency to include multiple patches on their interface regions. (B) The hub proteins have more interface mutations locating inside the patches. Only the proteins having at least one patch on their interface regions are shown.

### Patient-specific sub-networks inferred from mutation profiles cluster tumors based on pathway similarities

Given that disease-associated mutations significantly alter the interaction networks, we applied a network modeling approach to reveal the interactions from the mutation profile of each patient.

#### Patient-specific network reconstruction

We used Omics Integrator software [43] to reconstruct the mutation sub-network for each patient. Omics Integrator searches for the optimal network that connects the mutated proteins either directly or by adding intermediate nodes through high probability protein-protein interactions. In our setup, each protein having at least one missense mutation in the patient is added to the list as the base for network reconstruction and weighted with the number of mutations in that protein. We used the probability weighted protein-protein interactions in iRefWeb [44] as the reference interactome. Omics Integrator integrates the given list of protein hits with the interactome to find a parsimonious sub-network that connects the given proteins either directly or by adding intermediate proteins. The intermediate proteins are important in a sense that they connect the mutated proteins and behave as a complementing component of the pathways. Additionally, Omics Integrator has a unique feature to avoid the bias of the dominance of well-studied proteins or hubs in the final network. We used both hub-penalizing parameters set to reveal more specific pathways and non-penalizing parameters set to reveal canonical pathways for a better comparison of the patient-specific networks. Three different values of the scaling factor of hub proteins (μ parameter, described in Methods) are used for this purpose and resulting optimal networks were merged. As a result, a sub-network was found for each patient. In total 290 networks were reconstructed. The reconstructed networks consist of both mutated proteins in that patient and also many intermediate proteins those are connecting mutated proteins with high confidence edges. In this way, beyond the mutated proteins we are able to analyze how these proteins are affecting the rest of the sub-network.

#### Comparison and clustering of the patient-specific sub-networks

A comparison of the sub-networks is important to understand the commonalities and differences across the patients. In general, directly comparing the presence of the proteins and their interactions across the patients is straightforward. However, many cross talking pathways are present in these reconstructed networks and revealing those pathways is very important for a deep comparison. Therefore, we first applied an enrichment analysis of known pathways deposited in KEGG [45] to the reconstructed patient-specific pathways using Webgestalt [46]. The overrepresented pathways were retrieved based on a cutoff value 0.1 for FDR. To focus only on pathways that are not assigned to a disease we eliminated infections, cancers and addiction pathways. At the end, we came up with a union of 123 pathways. Among these, mTOR signaling, Jak-Stat and Ras signaling pathways have tendency to act together while TNF signaling and Nfkb signaling pathways show presence in another set of patients. This result guided us toward the classification of the networks based on the pathway similarities and then integrating this classification with the structural information of the mutations to identify signature spatial clusters of groups. To demonstrate that network-guided analysis have additional merits in the comparison of mutation profiles against simply analyzing the list of mutated proteins, we performed the same enrichment analysis using only the mutated proteins. In this analysis, we piped only the list of mutated proteins to Webgestalt analysis tool and we obtained overrepresented pathways for each patient. The results show that only in 11 patients there is a significant enrichment and the number of enriched pathways are only 6 which include EGFR signaling pathway and Glioma pathway that do not allow any further analysis to compare the patients and group them based on their similarities. These results also show that network reconstruction from mutation profiles reveals the affected pathways more extensively.

We compared the reconstructed patient-specific networks according to the overrepresented pathways and applied non-negative matrix factorization (NMF) implemented in GenePattern 2.0 [47] to cluster patients based on network inferred pathway similarities. As a result, we found four groups of GBM patients according to the mutation inferred pathway similarities. Many pathways are significantly enriched in each patient and while some pathways are significantly active in a group of patients those pathways are not present in the rest. In Fig 7A, the consensus heatmap is shown that in each group, the patients are consistently clustered together based on their pathway similarities. The resulting four groups contain 51, 108, 49 and 51 patients, respectively. Next, we merged the patient-specific networks of each identified group to find the patient-group-specific networks. As a result, we came up with four patient-group-specific consensus networks. We compared the four GBM subtypes (classical, neural, preneural, mesenchymal) derived from transcriptomic data with our network-based grouping. Each of our groups looked like a mixture of these subtypes. Then, we applied a hypergeometric test to see if any transcriptomic subtype is enriched in our groups. We found that group II and group IV are enriched in classical subtype (p-val=0.005, p-val=0.01, respectively). While group III is mainly enriched in mesenchymal subtype (p-val= 4.88 e-10), preneural and neural subtypes are also significantly present in group III (p-val=0.00014, p-val=0.025).

**Fig 7.**
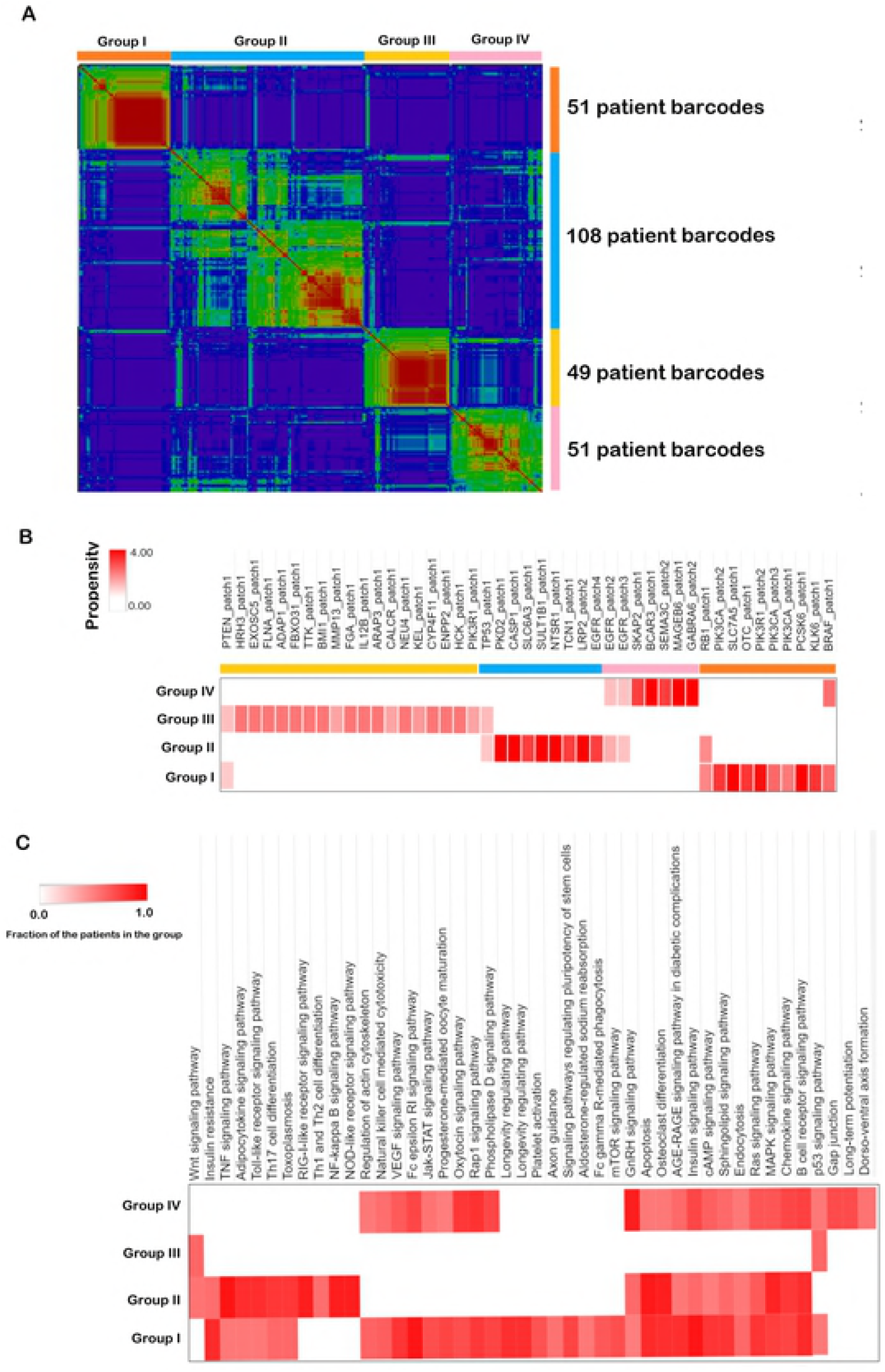
Classification of the tumors based on pathway similarities. (A) Consensus clustering of the network inferred disease signatures. (B) Propensities of each 3D mutation clusters in the group of patients. (C) The list of dominant pathways in each group of patients. Heatmaps are generated by Morpheus [48].

#### Identification of the marker 3D patches in network-inferred patient groups

Mutations in each patient are mostly located on the surface region (on average 65%) and the rest is in the core and interface regions. The same trend is also observed in the set of mutations of the four patient groups. Mutations on the surface and in the interface regions imply that they have a strong potential to alter the interaction network in the disease condition. Therefore, reconstructed networks are a rich source for a rigorous comparison across patients beyond the list of mutations. The arrangement of the mutations in 3D (patches) is also important to interpret with the network information. Our patient groups are based on the similarities of the networks that are inferred from patient-specific mutation profiles. The next question is whether any 3D mutation patches have a tendency to represent a patient group. To address this, we calculated the propensity of each 3D patch in each patient group (described in Methods). As a result we found that 45 patches have a tendency to be present in one or two patient groups. According to our results, the PIK3R1 patch, multiple patches of PIK3CA and BRAF have a strong tendency to be present in the group I (Fig 7B). While BRAF, RB1, PIK3CA, PIK3R1 patches are prone to the group I; TP53, CASP1, EGFR patches have bias to be in the group II. The mutations in hub proteins are organized into multiple patches. For example, EGFR has 5 patches, TP53 and PTEN have two patches. These patches are located in different groups. While one patch of PIK3R1 is present in the group I and the other is present in group IV. Patch 4 of EGFR is biased to be in group II, but Patch 2 and 3 are shared between groups II and IV. One of the very well characterized biomarkers in GBM is BRAF. BRAF mutations V600E and G596D form a 3D patch in group I and group IV.

Not only the patches but also the singleton mutations have a tendency to be present in some groups. IDH1 mutation is in group I (5 out of 12 patients with IDH1 mutations are in group I). Although mTOR is mutated only in one patient in group I, it is present in 14 other patients’ networks to connect mutated proteins and as a component of mTOR signaling pathway which is predominantly accumulated in group I (Fig 7C).

Interface mutations affect 323 interactions in group I, 256 interactions in group II, 380 interactions in group III and 200 interactions in group IV patients. In total, interface mutations are in 958 interactions in patient networks. Out of 290 patients, the interaction between EGFR and STUB1 proteins is the most affected interaction. There is at least one mutation in 31 patients at the binding region of EGFR and STUB1 proteins. It has been shown that STUB1 is an ubiquitination regulator of EGFR in cancer cells and a change in this interaction may increase the expression level of EGFR [49]. EGFR-STUB1 interaction is present in 17 patients in the group I.

We illustrated a sample merged network of the group I in Fig 8 where the interactions present in at least 3 tumors are drawn. The pie chart that is shown in each node represents the ratio of being mutated (red portion) or not (blue portion). The size of the nodes represents the frequency of that node to be present in group I. The thickness of the edges also represents the frequency of that edge to be present in group I. It is important to note that in this network there are intermediate proteins that connect mutated ones although they are not mutated. CTTNB1 is an example of an intermediate protein that is not mutated in any patient in the group I, but it connects many mutated proteins including CDH1, PTEN.

**Fig 8.**
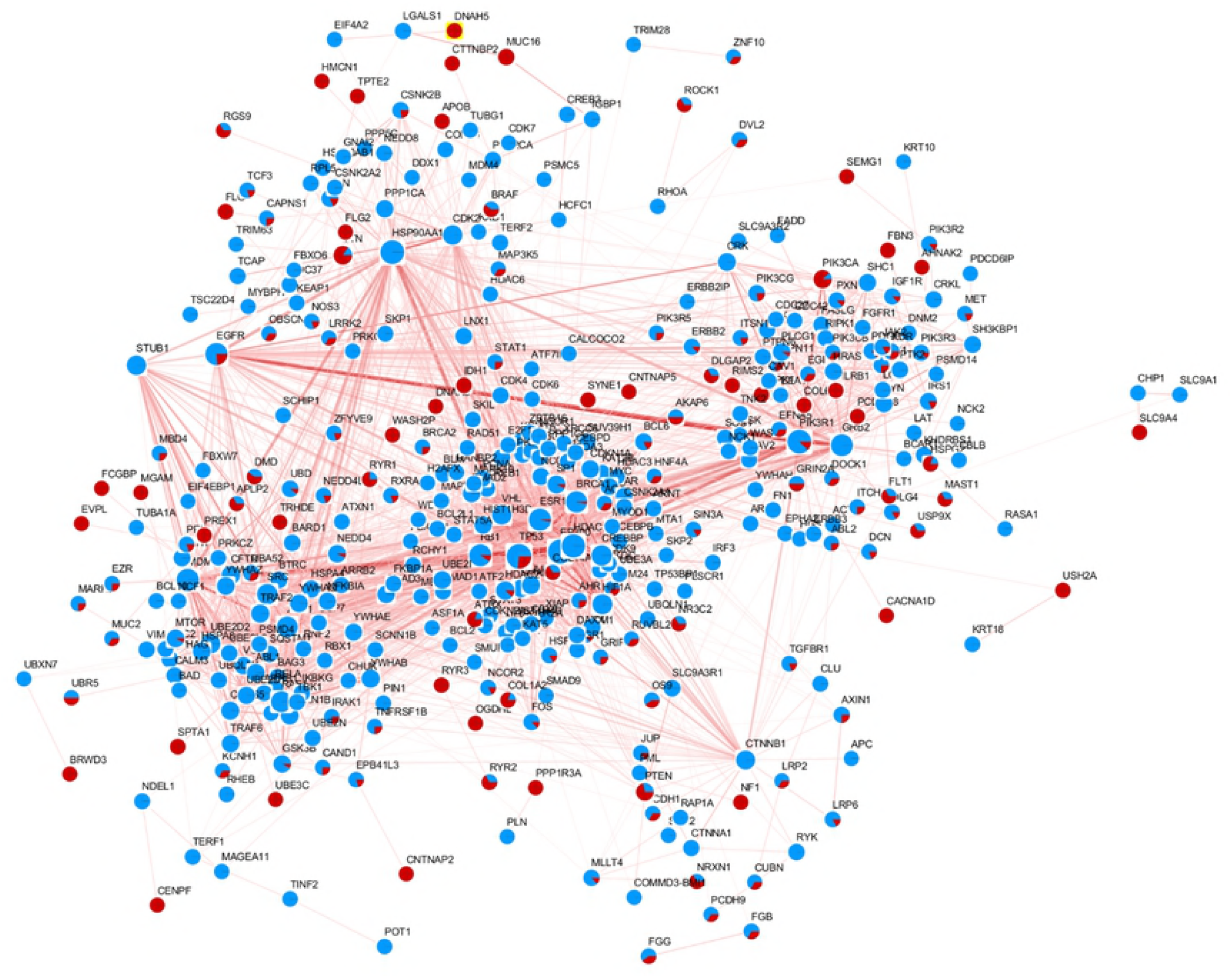
The merged network of group I. The edges that are present in at least three patients in Group I are drawn in this network. The nodes are labelled with a pie chart colored in red and/or blue color. The fraction of the red color represents the count of being a mutated protein in the patient network. The fraction of the blue color represents the count of being an intermediate protein connecting mutated ones in the patient network. Cytoscape is used for network visualization [50].

There are in total 1076 proteins in the union set of nodes in each network group of patients where the merged network of patient groups contain the edges that are present in at least three patients. Among them, 162 proteins are common in all groups, 584 proteins are present in only one patient group and 330 proteins are present in two or three patient groups. We ranked the proteins in each merged network of patient groups based on the node centrality. Among the top 20 most central proteins in each patient groups, EGFR, TP53, SRC and 10 more proteins are common across all groups. These proteins are previously defined as the “frequent flyers” that are the components of the many canonical signaling pathways and not discriminative across different networks [43]. However, there are also some central proteins that are specific to a patient group; such as IKBKG, which is among the top 20 central proteins only in group II.

## Conclusion

Glioblastoma (GBM) is a very heterogeneous disease. The set of GBM mutations is not discriminative to classify the progress and subtype of the disease. Here, we follow a systems level approach in which we integrate the mutations in GBM tumors with the protein structures/interfaces and reconstruct the patient specific networks. Our results show that out of 14644 mutations, 4392 mutations have structural information and 13% of these mutations are spatially grouped into patches while most of the mutations are spatially distant to other mutations, namely, are singletons. Despite a small portion of all mutations, 3D patches decrease this heterogeneity.

Mutated residues are mostly populated over the surface region and their functional effect is less severe than the rest (interface and core). Although a small portion of all mutations are located in the core region; they affect the function more severely. Mutations in the core region have a tendency to preserve their hydrophobicity. In hub proteins, mutations are organized into very large patches that connects mutations in multiple binding sites through the core of the protein. Additionally, hub proteins are adapted to have multiple 3D patches. The adaptation of hub proteins to have multiple interaction surfaces is also apparent in the 3D organization of their mutations according to our results. Our results suggest that hub proteins’ patch mutations are more disease causing, whereas other proteins’ singleton mutations are more disease causing. Interface mutations are divided into two groups that are shared and multiface. Some interface residues in the same protein affect more than one interaction because of the repeatedly usage of the same face with different partners that we call “shared” mutations. In some proteins mutations are located in more than one binding site, still affects multiple interactions but through different faces that we call “multi-face” mutations.

When we analyzed the patient specific networks and the consensus network of these patients, we observed that although mutation profiles are very heterogeneous across patients and their pathway level representation is very limited, the network-based analysis groups the patients better, reveals predominant pathways in each group. Additionally, the network-based similarity analysis shows that each group of patients carries a set of signature 3D mutation patches. For example, BRAF, RB1, PIK3CA, PIK3R1 patches are frequent in the group I, TP53, CASP1, EGFR patches are found in the group II. Beyond the list of mutations, network-guided analysis also reveals similarities across patients and overcomes the heterogeneity in mutation profiles by completing the interaction components that mutated proteins potentially affect. Overall, these results show that network-guided interpretation of mutations and their 3D organizations give a more diverse insight about their impact and can help in precision medicine.

## Methods

### Data collection and preprocessing

The missense mutations in Glioblastoma are retrieved from TCGA which has been published in [51] for 290 patients. First, all proteins that have at least one mutation is searched in PDB [31]. If a structure is not available, we made use of ModBase [32] homology models. The structural information of the protein interactions are collected from PDB, Interactome3D [6], PRISM [37] and Interactome Insider [13]. PDB deposits protein complexes that are crystallized together. Interactome3D predicts the protein complexes through structure and domain similarity with a template structure. PRISM uses known interfaces to predict new protein interactions. We used the pre-runned PRISM results for a subset of the proteome, not the whole proteome. The other source for structural protein interactions is the Interactome Insider. It produces the binding sites on each partner of the protein interaction. Different than PRISM and Interactome3D, it does not give the structure or the pose of the predicted protein complex. We also downloaded the human proteome from UniProt [52] for cross-referencing from one data source to another. Additionally, the residue positions in sequence are not consistent with the residue positions in protein structures. A PDB entry or a homology model of a UniProt sequence may represent only a fragment of the given protein and the residue numbering may not be exactly the same with the sequence positions. Therefore, we performed UniProt sequence to the sequence in the protein structure alignment to find the exact position of each residue.

Finally, the confidence weighted interactome deposited in iRefWeb has been downloaded for the reconstruction of patient specific sub-networks inferred from mutation profile of each patient.

### Identification of the spatial clusters

Each protein structure and protein complex were converted in a residue-residue interaction network. If any atom in a residue is in a close proximity to any atom in another residue, then these two residues were considered to be interacting. The proximity is defined as the distance less than 5Å between any atoms. At the end for each structure we constructed a residue contact graph R(v, e) where v is the set of residues and e is the set of edges between these residues. To identify the spatial clusters, we searched for all shortest paths between each mutated residue pairs with a length less than 3 which means that if two mutated residues are either connected directly or only one residue is in between them. Then all the extracted shortest paths merged to come up with a subgraph P(v’,e’) representing one spatial cluster, namely “patch” where v’⊂v and e’⊂e. Mutations that are not assigned to a patch are labelled as singleton that means these residues are distant to other mutations in the same protein.

### Identification of protein regions and the effect of the mutations

Proteins can be divided into three regions; namely, core, surface and interface regions. The conventional approach for identification of these regions are calculating solvent accessible surface areas of each residue in the protein. FreeSASA [53] is a software designed for calculation of solvent accessible surface area both at residue level and at molecule level. In general, if the relative solvent accessible surface area of a residue in its monomer state is greater than or equal to 15%, then this residue is labelled as the surface residue. Interface residues that are collected from structural interactome are excluded from the surface residue set. The rest is identified as core residues.

To calculate the effect of mutations if they are damaging or neutral, we used EVmutation and PolyPhen-2 web servers. The EVmutation data is provided for a limited number of proteins in text format where each position in a UniProt entry is substituted to the remaining 19 amino acid and the damage score is calculated. The more negative values of the calculated score means the more damaging mutation. We used this data to measure the likelihood of our mutations and we took the score of the displacement that occurred in each mutation and compared it with the average score of all other displacement probabilities. For this purpose, for each matched UniProt entries of each mutation, we retrieved a score representing the damage.

### Sub-network reconstruction for each patient

Omics Integrator software was used to reconstruct patient-specific sub-networks. Given a reference graph G(V, E, w) where V is the node set {v|v εV} and {e|e ε E} is the edge set and w is the edge weights, the Forest module of Omics Integrator solves the prize-collecting Steiner forest problem for a given set of nodes with predefined prizes and returns an optimal network that covers a subset of the given proteins having prize and additional intermediate proteins that connects the proteins having prize. Prizes were given as the number of mutations per protein for each patient in our case. Therefore, the prize list composed of the proteins having at least one mutation. We used the iRefWeb v8.0 as the weighted reference interactome in our modeling. The parameter set ω= 10.0, depth (D) = 10 and β = 5 was used for the reconstruction. ω parameter was used for tuning the number of trees in the final network, depth is the number of edges from the root to the leaf nodes and β is a scaling factor to force more prize nodes to enter the final network. Finally, μ is another scaling factor to tune the dominance of hub proteins in the final network. We used three μ values (0.0, 0.005. 0.01) to recover the canonical pathways and more specific ones and merged the node and edge set of the reconstructed networks to come up with a single network for each patient.

### Classification of the patients into groups based on reconstructed networks

Sub-networks of each patient were analyzed with Webgestalt tool to retrieve the overrepresented KEGG pathways in each network. Pathways were assumed to be enriched in the sub-network if the FDR is less than 0.1. In the resulting list of pathways, we eliminated disease pathways including infections, cancer, addiction related pathways. Then, we prepared a matrix where rows are union set of enriched pathways, columns are patient barcodes and the entries are the enrichment score (ES) of a pathway in the corresponding barcode’s sub-network. If the pathway is not enriched than 0 is inserted to that entry. This matrix is piped to the consensus clustering based on non-negative matrix factorization (NMF) approach implemented in GenePattern 2.0 with the k parameter from 2 to 5.

The identified groups were searched for if any identified spatial patch has tendency to represent a group. We calculated the propensity of group i for having patch j, P_ij_. Equation (1) was used where x_i_j is the barcode set in the group i having at least one mutation in patch j, n_i_ is the number of barcodes in group i, X_j_ is the number of barcodes having at least one mutation in patch j and N is the total number of barcodes.

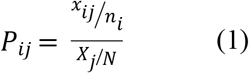

## Acknowledgements

N.T. acknowledges the support from the Career Development Program of TUBITAK under the project number 117E192 and UNESCO-L’Oreal National for Women in Science Fellowship. Authors thank the Science Academy, Turkey.

## Supporting information

**S1 File. Interface mutations and the interactions that are related for each mutation.** For each 791 interface mutations, corresponding interactions that are related by the mutation are listed in tab separeted format.

